# SPECTRE: a Suite of PhylogEnetiC Tools for Reticulate Evolution

**DOI:** 10.1101/169177

**Authors:** Sarah Bastkowski, Daniel Mapleson, Andreas Spillner, Taoyang Wu, Monika Balvočiūtė, Vincent Moulton

## Abstract

**Summary:** Split-networks are a generalization of phylogenetic trees that have proven to be a powerful tool in phylogenetics. Various ways have been developed for computing such networks, including split-decomposition, NeighborNet, QNet and FlatNJ. Some of these approaches are implemented in the user-friendly SplitsTree software package. However, to give the user the option to adjust and extend these approaches and to facilitate their integration into analysis pipelines, there is a need for robust, open-source implementations of associated data structures and algorithms. Here we present SPECTRE, a readily available, open-source library of data structures written in Java, that comes complete with new implementations of several pre-published algorithms and a basic interactive graphical interface for visualizing planar split networks. SPECTRE also supports the use of longer running algorithms by providing command line interfaces, which can be executed on servers or in High Performance Computing (HPC) environments.

**Availability:** Full source code is available under the GPLv3 license at: https://github.com/maplesond/SPECTRE

SPECTRE’s core library is available from Maven Central at: https://mvnrepository.com/artifactuk.ac.uea.cmp.spectre/core

Documentation is available at: http://spectre-suite-of-phylogenetic-tools-for-reticulate-evolution.readthedocs.io/en/latest/

**Supplementary Information (SI):** Supplementary information is available at Bioinformatics online.

## 1 INTRODUCTION

Split-networks are a generalization of phylogenetic trees that are commonly used to analyze reticulate evolution in organisms such as plants, bacteria and viruses (see Figure 1 for an example). They provide a snapshot of the data and can be used to display conflicting signals. Examples of algorithms for computing such networks include split-decomposition (Bandelt and Dress, 1992), Neighbor-Net (Bryant and Moulton, 2004), QNet (Grünewald *et al.*, 2007), SuperQ (Grünewald *et al.*, 2013) and FlatNJ (Balvočiūtė *et al.*, 2014). A comprehensive overview of split-networks can be found in (Huson and Bryant, 2006).

Currently the main program available for computing split-networks is the user-friendly SplitsTree program (Huson and Bryant, 2006). In addition, various methods for computing split-networks such as some of those mentioned above have been implemented and released as stand alone applications. Implementing data structures capable of representing the mathematical structures used to describe and compute split networks is not a trivial undertaking and existing software either is closed source or have their data structures and algorithms tightly integrated with their host tool, so are not easily reusable. There are therefore currently few options for developers wishing to create or extend their own tools based on these concepts other than to start from scratch. Hence there is a need for a robust and flexible open-source library that provides core data structures and algorithms to facilitate development of new tools and algorithms.

## 2 SPECTRE

Here we present SPECTRE, a suite of tools for computing, modelling, and visualizing reticulate evolution based on split-networks. SPECTRE builds in part on existing open-source implementations of some of these tools, in particular for QNet, SuperQ and FlatNJ, integrating them into a unified and extendible library. The main tools available through SPECTRE are summarized below (for more details see section 1 of SI):

- NeighborNet rapidly constructs a circular split network from a distance matrix or a sequence alignment (Bryant and Moulton, 2004). NetMake implements variants of NeighborNet as described in (Levy and Pachter, 2011).
- SuperQ constructs a circular split network from a set of (partial) input trees (Grünewald *et al.*, 2013).
- FlatNJ constructs a flat split network from a multiple sequence alignment, weighted quartet data or location data (Balvočiūtė *et al.*, 2014).
- NetME produces a minimum evolution tree compatible with an existing circular split network (Bastkowski *et al.*, 2014).

**Fig. 1:**
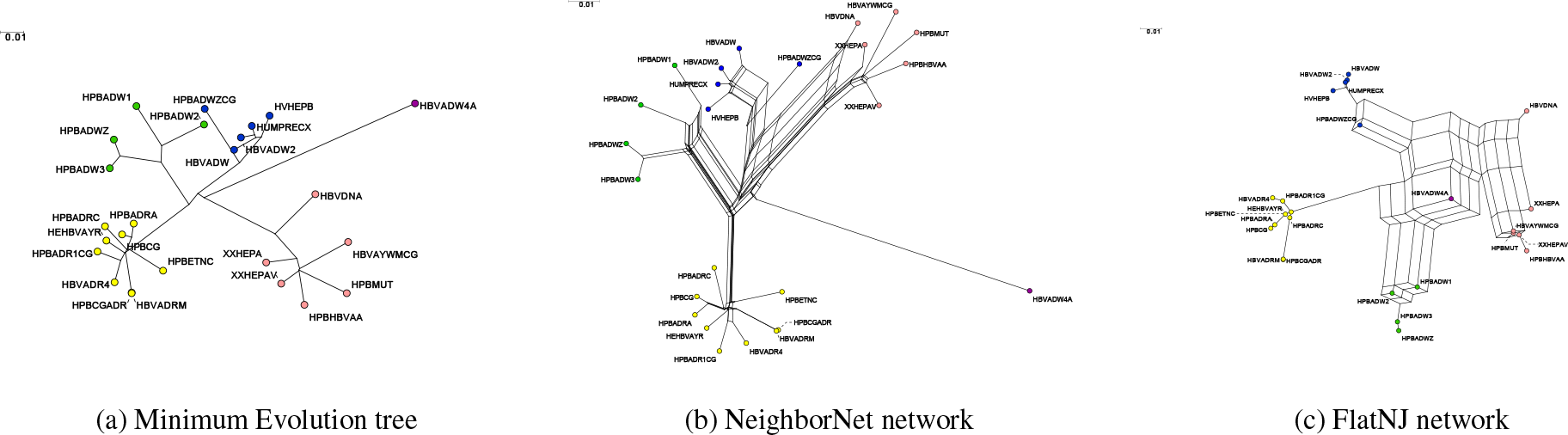
To illustrate some of SPECTREs functionality, we processed a data set analyzed in (Bollyky *et al.*, 1996) consisting of different Hepatitis B viruses (HBV). There are 5 different genomic groups and the phylogenetic analysis led to the result that HBVDNA is a recombinant with around half the genome coming from group A and half from group D. It also concluded that HPBADW1 is a recombinant of HPBADW2 (B) and HPBADWZCG (A), but with only a small insertion from HPBADWZCG into the Genome. (a) shows a minimum evolution tree constructed by NetME that is compatible with the split network constructed by NeighborNet, which is shown in (b). (c) shows the split network constructed by FlatNJ.

These tools are accessible to the user via graphical and command line interfaces. Apart from driving the tools, the interactive graphical interface can visualize planar split networks using the drawing algorithm in (Spillner *et al.*, 2012). The interface offers a number of basic functions for orientating the canvas (e.g. zoom, pan, flip and rotate), manipulating labels (size, color, location) and creating image files (PDF, EPS, SVG, PNG). The command line implementation enables bioinformaticians to integrate tools into pipelines. This works on desktop PCs, like SplitsTree, but is also designed so long running tools are executable on servers or high performance computing environments where displays are not available.

For developers wishing to reuse code and develop their own tools, SPECTRE provides a core library containing common data structures (e.g. splits, trees, networks, distances, quartets and multiple sequence alignments), algorithms (e.g. NeighborNet), and robust file parsers to process a range of input files (e.g. NEXUS, PHYLIP, Newick, Emboss, FastA); see section 2 of SI for more details. The library is available directly from Maven Central, giving developers direct access to the most recent version of the library and and providing a convenient way to integrate it into the processes for building their own projects.

## 3 CONCLUDING REMARKS

SPECTRE provides a collection of open-source tools and resources for modelling, understanding and visualizing reticulate evolution based on split networks. We believe that our software will both enable bioinformaticians to easily test and compare methods for inferring planar split networks and help computer scientists build their own methods for inferring phylogenetic networks by reusing our existing data structures and algorithms via the open-source library. Moreover, this also provides the option to easily add such new tools to the library making them readily available to other users.

## ACKNOWLEDGMENTS

We would like to thank Stephan Grünewald.

